# Forecasting seasonal influenza in the U.S.: A collaborative multi-year, multi-model assessment of forecast performance

**DOI:** 10.1101/397190

**Authors:** Nicholas G Reich, Logan Brooks, Spencer Fox, Sasikiran Kandula, Craig McGowan, Evan Moore, Dave Osthus, Evan Ray, Abhinav Tushar, Teresa Yamana, Matthew Biggerstaff, Michael A Johansson, Roni Rosenfeld, Jeffrey Shaman

## Abstract

Influenza infects an estimated 9 to 35 million individuals each year in the United States and is a contributing cause for between 12,000 and 56,000 deaths annually. Seasonal outbreaks of influenza are common in temperate regions of the world, with highest incidence typically occurring in colder and drier months of the year. Real-time forecasts of influenza transmission can inform public health response to outbreaks. We present the results of a multi-institution collaborative effort to standardize the collection and evaluation of forecasting models for influenza in the US for the 2010/2011 through 2016/2017 influenza seasons. For these seven seasons, we assembled weekly real-time forecasts of 7 targets of public health interest from 22 different models. We compared forecast accuracy of each model relative to a historical baseline seasonal average. Across all regions of the US, over half of the models showed consistently better performance than the historical baseline when forecasting incidence of influenza-like illness 1, 2 and 3 weeks ahead of available data and when forecasting the timing and magnitude of the seasonal peak. In some regions, delays in data reporting were strongly and negatively associated with forecast accuracy. More timely reporting and an improved overall accessibility to novel and traditional data sources are needed to improve forecasting accuracy and its integration with real-time public health decision-making.

## Introduction

Over the past 15 years, the number of published research articles on forecasting infectious diseases has tripled (Web of Science). This increased interest has been fueled in part by the promise of ‘big data’, that near real-time data streams of information ranging from large-scale population behavior [1] to microscopic changes in a pathogen [2] could lead to measurable improvements in how disease transmission is measured, forecasted, and controlled [3]. With the spectre of a global pandemic looming, improving infectious disease forecasting continues to be a central priority of global health preparedness efforts.[4, 5, 6]

Forecasts of infectious disease transmission can inform public health response to outbreaks. Accurate forecasts of the timing and spatial spread of infectious disease incidence can provide valuable information about where public health interventions can be targeted.[7] Decisions about hospital staffing, resource allocation, the timing of public health communication campaigns, and the implementation of interventions designed to disrupt disease transmission, such as vector control measures, can be informed by forecasts. In part due to the growing recognition of the importance of systematically integrating forecasting into public health outbreak response, large-scale collaborations have been used in forecasting applications to develop common data standards and facilitate comparisons across multiple models.[8, 9, 10, 11] By enabling a standardized comparison in a single application, these studies greatly improve our understanding of which models perform best in certain settings, of how results can best be disseminated and used by decision-makers, and of what the bottlenecks are in terms of improving forecasts.

While multi-model comparisons exist in the literature for single-outbreak performance [8, 11, 10], here we compare a consistent set of models over seven influenza seasons. To our knowledge, this is the first documented comparison of multiple real-time forecasting models from different teams across multiple seasons for any infectious disease application. Since each season has a unique dynamical structure, multi-season comparisons like this have great potential to improve our understanding of how models perform over the long term and which models may be reliable in the future.

Influenza is a respiratory viral infection that can cause mild or severe symptoms. In the US each year, influenza viruses infect an estimated 9 to 35 million individuals and cause between 12,000 and 56,000 deaths.[12] Influenza incidence typically exhibits a strong annual periodicity in the US (and in other global regions with temperate climates), often circulating widely during colder months (i.e., November through April). The social, biological, environmental, and demographic features that contribute to higher-than-usual incidence in a particular season are not fully understood, although contributing factors may include severity of the dominant influenza subtype[13], temperature and humidity[14], vaccine effectiveness[12], or timing of school vacations[15].

Starting in the 2013/2014 influenza season, the U.S. Centers for Disease Control and Prevention (CDC) has run the “Forecast the Influenza Season Collaborative Challenge” (a.k.a. FluSight) each influenza season, soliciting prospective, real-time weekly forecasts of regional-level weighted influenza-like illness (wILI) measures from teams across the world (Figure 1).[8, 10] The FluSight challenge focuses on forecasts of the weighted percentage of doctor’s office visits where the patient showed symptoms of an influenza-like illness in a particular region. Weighting is done by state population as the data are aggregated to the regional and national level. This wILI metric is a standard measure of seasonal flu activity, for which public data are available for the US back to the 1997/1998 influenza season (Figure 1A). The FluSight challenge forecasts are displayed together on a website in real-time and are evaluated for accuracy at the end of the season.[16] This effort has galvanized a community of scientists interested in forecasting, creating a testbed for improving both the technical understanding of how different forecast models perform and the integration of these models into decision-making.

**Figure 1:**
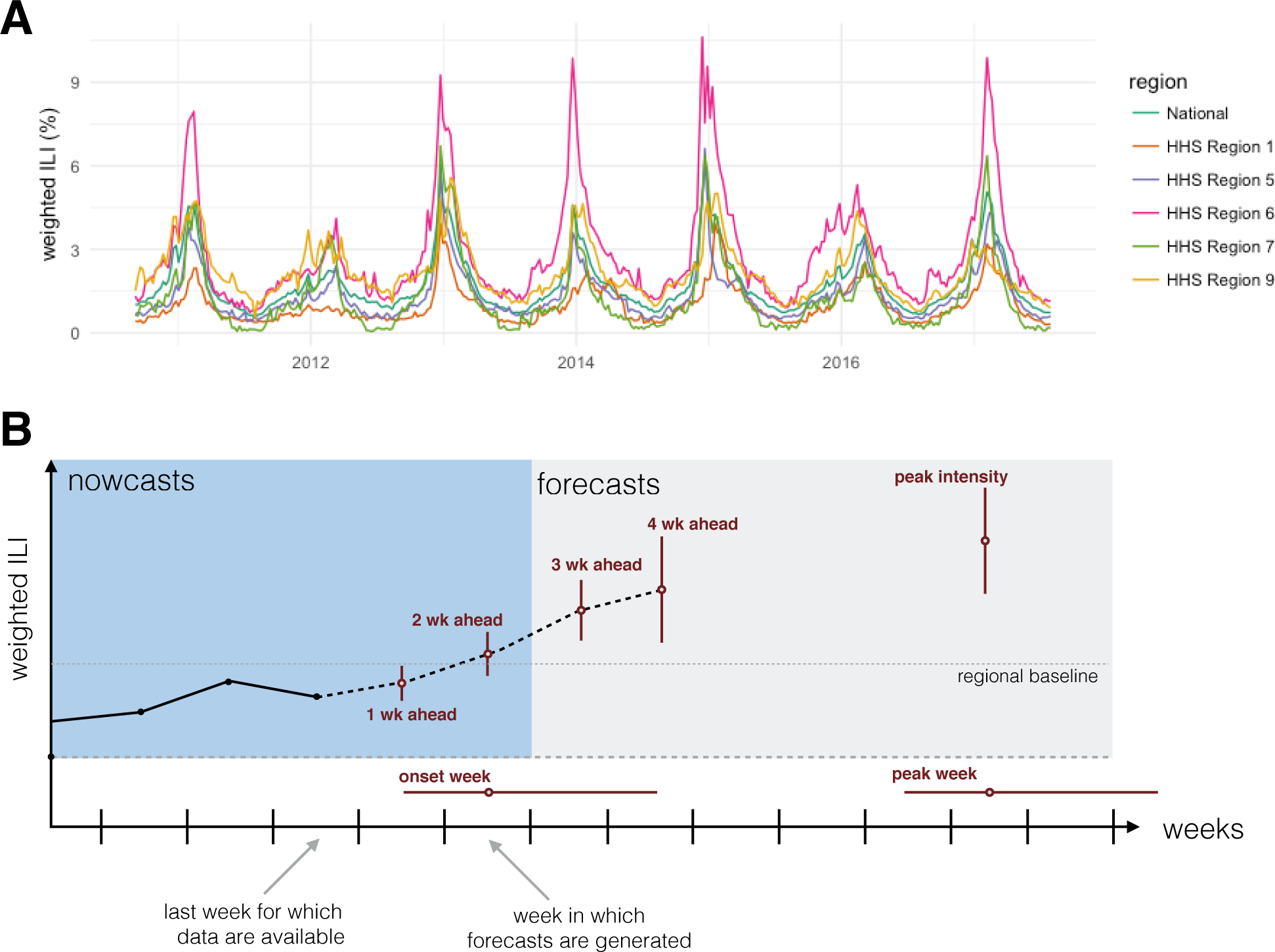
(A) Weighted influenza-like illness data downloaded from the CDC website from selected regions. The y-axis shows the weighted percentage of doctor’s office visits in which a patient presents with influenza-like illness for each week from September 2010 through July 2017, which is the time period for which the models presented in this paper made seasonal forecasts. (B) A diagram showing the anatomy of a single forecast. The seven forecasting targets are illustrated with a point estimate (dot) and interval (uncertainty bars). The five targets on the wILI scale are shown with uncertainty bars spanning the vertical wILI axis, while the two targets for a time-of-year outcome are illustrated with horizontal uncertainty bars along the temporal axis. The onset is defined relative to a region-and season-specific baseline wILI percentage defined by the CDC.[17] Arrows illustrate the timeline for a typical forecast for the CDC FluSight challenge, assuming that forecasts are generated or submitted to the CDC using the most recent reported data. These data include the first reported observations of wILI% from two weeks prior. Therefore, 1 and 2 week-ahead forecasts are referred to as nowcasts, i.e., at or before the current time. Similarly, 3 and 4 week-ahead forecasts are forecasts, or estimates about events in the future.

Building on the structure of the FluSight challenges (and those of other collaborative forecasting efforts [9, 11]), a subset of FluSight participants formed a consortium in early 2017 to facilitate direct comparison and fusion of modeling approaches. Our work brings together 22 models from five different institutions: Carnegie Mellon University, Columbia University, Los Alamos National Laboratory, University of Massachusetts-Amherst, and University of Texas-Austin (Table 1). In this paper, we provide a detailed analysis of the performance of these different models in forecasting the targets defined by the CDC FluSight challenge organizers (Figure 1B). Drawing on the different expertise of the five teams allows us to make fine-grained and standardized comparisons of distinct approaches to disease incidence forecasting that use different data sources and modeling frameworks.

**Table 1:**
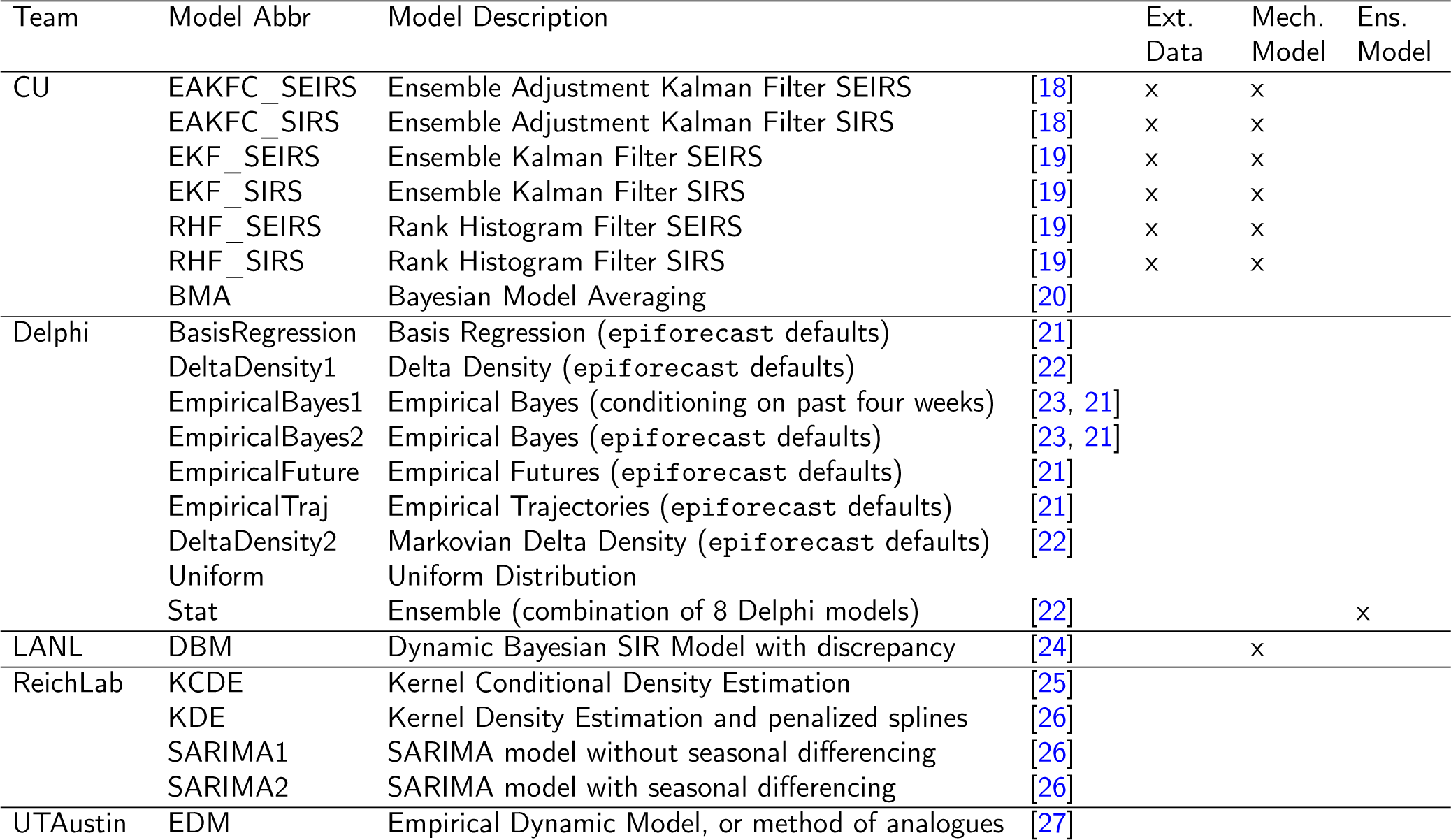
List of models, with key characteristics. Team abbreviations are translated as: CU = Columbia University, Delphi = Carnegie Mellon, LANL = Los Alamos National Laboratories, ReichLab = University of Massachusetts Amherst, UTAustin = University of Texas Austin. The ‘Ext data’ column notes models that use data external to the ILINet data from CDC. The ‘Mech. model’ column notes models that rely to some extent on an mechanistic or compartmental model formulation. The ‘Ens. model’ column notes models that are ensemble models.

In addition to analyzing comparative model performance over multiple seasons, this work identifies key bottlenecks that limit the accuracy and generalizability of current forecasting efforts. Specifically, we present quantitative analyses of the impact that incomplete or partial case reporting has on forecast accuracy. Additionally, we assess whether purely statistical models show similar performance to models that consider explicit mechanistic models of disease transmission. Overall, this work shows strong evidence that carefully crafted forecasting models for region-level influenza in the US consistently outperformed a historical baseline model for targets of particular public health interest.

## Results

### Performance in forecasting week-ahead incidence

Influenza forecasts have been evaluated by the CDC primarily using a variation of the log-score, a measure that evaluates both the precision and accuracy of a forecast.[28] Consistent with the primary evaluation performed by the CDC, we used a modified form of the log-score to evaluate forecasts (see Methods). The reported scores are aggregated into an average on the log scale and then exponentiated so the reported scores can be interpreted as the (geometric) average probability assigned to the eventually observed value of each target by a particular model. Therefore, higher scores reflect more accurate forecasts. As a common reference point, we compare all models to a historical baseline model, ReichLab-KDE, which forecasts the same historical average every week within a season and does not update based on recently observed data.

Average score for all of the short term forecasts (1 through 4 week-ahead targets) varied substantially across models and regions (Figure 2). The model with the highest average score for the week-ahead targets across all regions and seasons was CU-EKF_SIRS. This model achieved a region-specific average forecast score for week-ahead targets between 0.32 and 0.55. As a comparison, the historical baseline model ReichLab-KDE achieved between 0.12 and 0.37 average score for all week-ahead targets.

**Figure 2:**
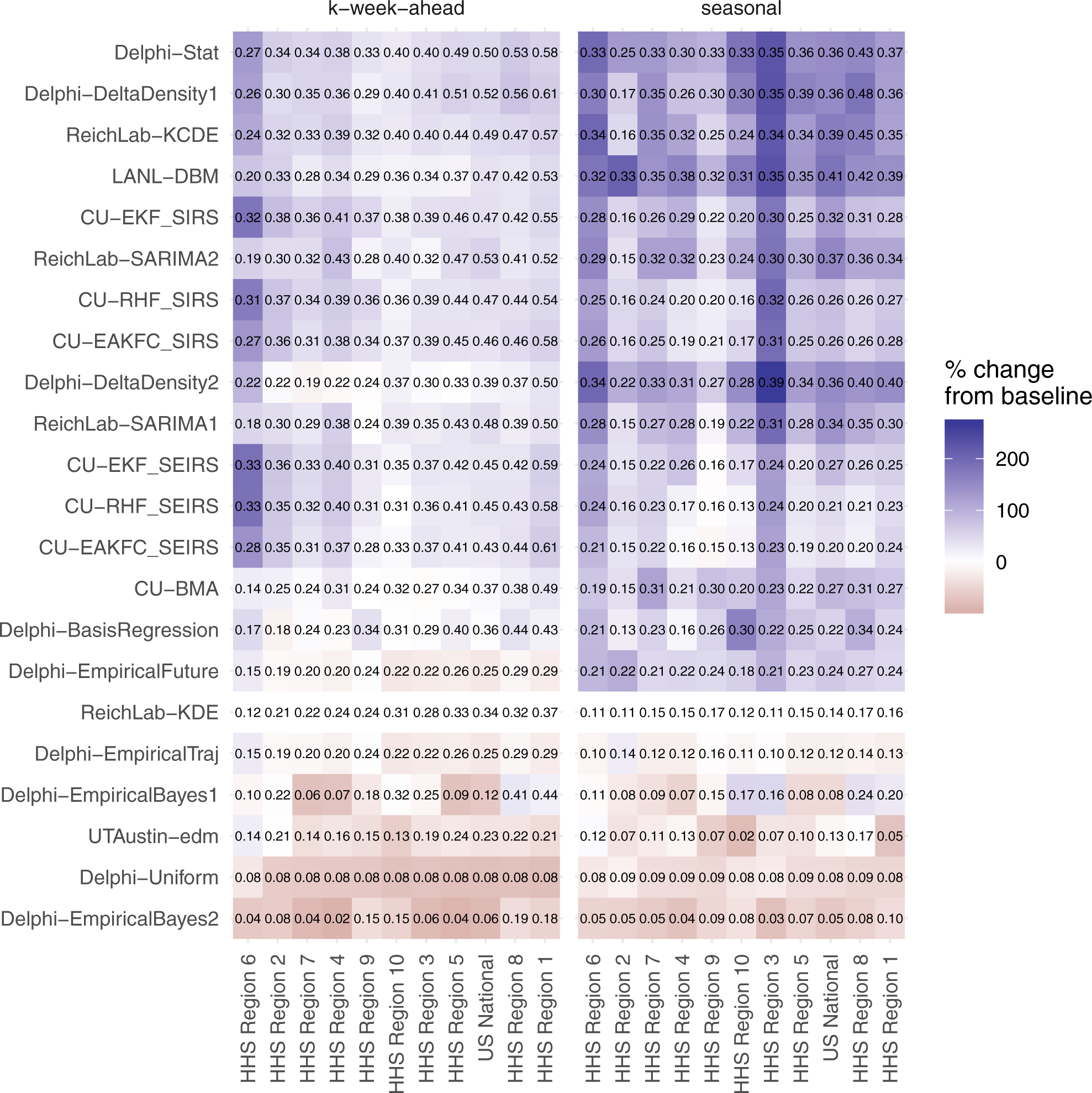
Average forecast score by model region and target-type, averaged over weeks and seasons. The text within the grid shows the score itself. The white midpoint of the color scale is set to be the target-specific average of the historical baseline model, ReichLab-KDE, with darker blue colors representing models that have better scores than the baseline and darker red scores representing models that have worse scores than the baseline. The models are sorted in descending order from most accurate (top) to least accurate (bottom) and regions are sorted from high scores (right) to low scores (left).

Models were more consistently able to forecast week-ahead wILI in some regions than in others. Predictability for a target can be broken down into two components. First, what is the baseline score that a model derived solely from historical averages can achieve? Second, by using alternate modeling approaches, how much more accuracy can be achieved beyond this historical baseline? Looking at results across all models, HHS Region 1 was the most predictable and HHS Region 6 was the least predictable (Figure 2).

The models presented show substantial improvements in accuracy compared to forecasts from the historical baseline model in all regions of the US. Results that follow are based on summaries from those models that on average showed higher forecast score than the historical baseline model. HHS Region 1 showed the best overall week-ahead predictability of any region. Here, the models showed an average forecast score of 0.54 for week-ahead targets (Figure 3A). This means that in a typical season these models assigned an average of 0.54 probability to the accurate wILI percentages. This resulted from having the highest score from the baseline model (0.37) and having the largest improvement upon baseline predictions (0.17) from the other models (Figure 3B). In HHS Region 6 the average week-ahead score was 0.24. While HHS Region 6 showed the lowest baseline model score of any region (0.12), it also showed the second highest improvement (0.12) upon baseline predictions (Figure 3B).

**Figure 3:**
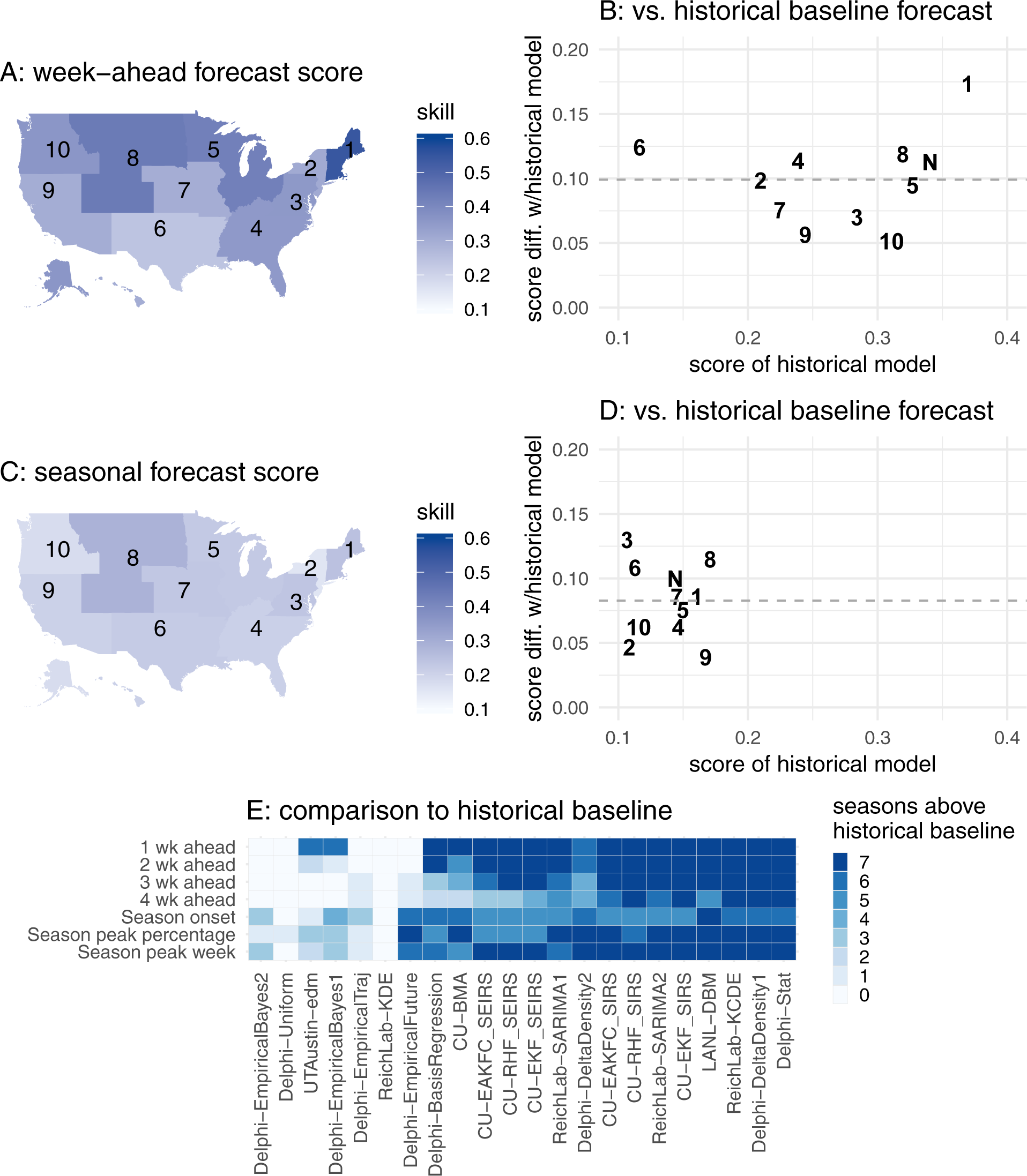
Absolute and relative forecast performance for week-ahead (top row) and seasonal (middle row) targets, summarized across all models that on average performed better than the historical baseline. Panels A & C show maps of the U.S. that illustrate spatial patterns of average forecast accuracy for week-ahead (A) and seasonal (C) targets. Color shading indicates average forecast score for this model subset. Panels B & D compare historical baseline model score (x-axis) with the average score (y-axis, horizontal dashed line at average across regions) with one point for each region. For example, a y-value of 0.1 indicates that the models on average assigned 10% more probability to the eventually observed value than the historical baseline model. The digits in the plot refer to the corresponding HHS Region number, with N indicating the US National region. Panel E shows the number of seasons each model had average performance above the historical baseline.

Forecast score declined as the target moved further into the future relative to the most recent observation. Over half of the models outperformed the historical baseline model in making 1-week ahead forecasts, as 15 of 22 models outperformed the historical baseline in at least 6 of the 7 seasons. However, only 7 out of 22 models outperformed the historical baseline in at least 6 seasons when making 4-week ahead forecasts. For the model with highest forecast score across all four week-ahead targets (CU-EKF_SIRS), the average scores across regions and seasons for 1 through 4 week-ahead forecasts were 0.55, 0.44, 0.36, and 0.31. This mirrored an overall decline in score observed across most models. Only in HHS Region 1 were the forecast scores from the CU-EKF_SIRS model for both the “nowcast” targets (1 and 2 weeks ahead) above 0.5.

### Performance in forecasting seasonal targets

Overall, forecast score was lower for seasonal targets than for week-ahead targets, although the models showed greater relative improvement compared to the baseline model (Figure 2). The historical baseline model achieved an overall forecast score of 0.14. The best single model across all seasonal targets was LANL-DBM with an overall forecast score of 0.36, more than a two-fold increase in score over the baseline.

Of the three seasonal targets, models showed the lowest average score in forecasting season onset, with an overall average score of 0.15. Due to the variable timing of season onset, different numbers of weeks were included in the final scoring for each region-season (see Methods). Of the 77 region-seasons evaluated, 9 had no onset, i.e., the wILI did not remain above a fixed region-specific threshold of influenza activity for three or more weeks (see Methods for details). The best model for onset was LANL-DBM, with an overall average score of 0.33 and region-season-specific scores for onset that ranged from 0.03 to 0.81. The historical baseline model showed an average score of 0.11 in forecasting onset. Overall, 8 out of 22 models (36%) had better overall score for onset in at least 6 out of the 7 seasons evaluated (Figure 3E).

Models showed an overall average score of 0.23 in forecasting peak week. The best model for peak week was ReichLab-KCDE, with an overall average score of 0.35. Region-and season-specific forecast score from this model for peak week ranged from 0.01 to 0.67. The historical baseline model showed 0.17 score in forecasting peak week. Overall, 15 out of 22 models (68%) had better overall score for peak week in at least 6 out of the 7 seasons evaluated (Figure 3E).

Models showed an overall average score of 0.20 in forecasting peak intensity. The best model for peak intensity was LANL-DBM, with overall average score of 0.38. Region-and season-specific forecast scores from this model for peak intensity ranged from 0.13 to 0.61. The historical baseline model showed 0.13 score in forecasting peak intensity. Overall, 12 out of 22 models (55%) had better overall score in at least 6 out of the 7 seasons evaluated (Figure 3E).

### Comparing models’ forecasting performance by season

Averaging across all targets and locations, forecast scores varied widely by model and season (Figure 4). The historical baseline model (ReichLab-KDE) showed an average seasonal score of 0.20, meaning that in a typical season, across all targets and locations, this model assigned on average 0.20 probability to the eventually observed value. The model with the highest average seasonal forecast score (Delphi-Stat) and lowest (Delphi-EmpiricalBayes2) had scores of 0.37 and 0.07, respectively. Of the 22 models, 16 models (73%) showed higher average seasonal forecast score than the historical average. Season-to-season variation was substantial, with 10 models having at least one season with greater average forecast score than the Delphi-Stat model did.

**Figure 4:**
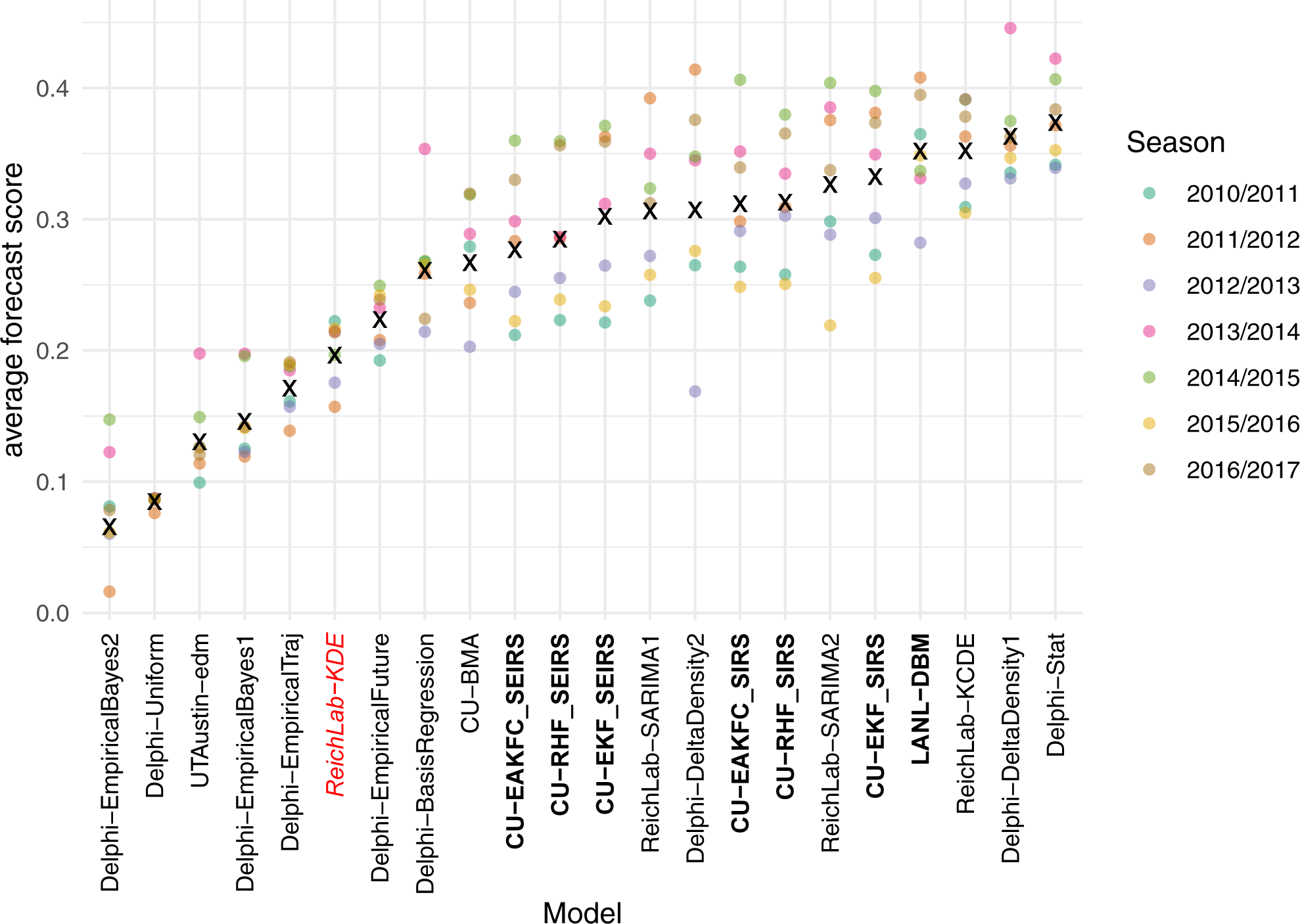
Average forecast score, aggregated across targets, regions, and weeks, plotted separately for each model and season. Models are sorted from lowest scores (left) to highest scores (right). Higher scores indicate better performance. Dots show average scores across all targets, regions, and weeks within a given season. The ‘x’ marks the geometric mean of the seven seasons. The names of compartmental models are shown in bold face. The ReichLab-KDE model (italicized red font) is considered the historical baseline model.

The six top-performing models utilized a range of methodologies, highlighting that very different approaches can result in very similar overall performance. The overall best model was an ensemble model (Delphi-Stat) that used a weighted combination of other models from the Delphi group. Both the ReichLab-KCDE and the Delphi-DeltaDensity1 model utilized kernel conditional density estimation, a non-parametric statistical methodology that is a distribution-based variation on nearest-neighbors regression. These models used different implementations and different input variables, but showed similarly strong performance across all seasons. The UTAustin-edm and Delphi-DeltaDensity2 models also used variants of nearest-neighbors regression, although overall scores for these models were not consistent, indicating that implementation details and/or input variables can impact the performance of this approach. The LANL-DBM and CU-EKF_SIRS models both rely on a compartmental model of influenza transmission; however the methodologies used to fit and forecast were different for these approaches. The ReichLab-SARIMA2 model used a classical statistical time-series model, the seasonal auto-regressive integrated moving average model (SARIMA), to fit and generate forecasts. Interestingly, several pairs of models, although having strongly contrasting methodological approaches, showed similar overall performance; e.g., CU-EKF_SIRS and ReichLab-SARIMA2, LANL-DBM and ReichLab-KCDE.

### Comparison between statistical and compartmental models

On the whole, statistical models achieved similar or slightly higher scores as compartmental models when forecasting both week-ahead and seasonal targets, although the differences were small and of minimal practical significance. Using the best three overall models from each category, we computed the average forecast score for each combination of region, season, and target (Table 2). For all targets, except 1 week-ahead forecasts and peak intensity, the difference in model score was slight, and never greater than 0.02. For 1-week-ahead forecasts, the statistical models had slightly higher scores on average than mechanistic models (0.06, on the probability scale). We note that the 1 week-ahead forecasts from the compartmental models from the CU team are driven by a statistical “nowcast” model that uses data from the Google Search application programming interface (API).[29] Therefore, the CU models were not counted as mechanistic models for 1 week-ahead forecasts. For peak percentage forecasts, the statistical models had slightly higher scores on average than mechanistic models (0.05).

**Table 2:** Comparison of the top three statistical models (Delphi-DeltaDensity1, ReichLab-KCDE, ReichLab-SARIMA2) and the top three compartmental models, (LANL-DBM, CU-EKF_SIRS, CU-RHF_SIRS) based on best average region-season forecast score. The difference column represents the difference in the average probability assigned to the eventual outcome for the target in each row. Positive values indicate the top statistical models showed higher average score than the top compartmental models.

| target | stat. model score | compartmental model score | difference |
| --- | --- | --- | --- |
| 1 wk ahead | 0.49 | 0.43 | 0.06 |
| 2 wk ahead | 0.40 | 0.41 | -0.01 |
| 3 wk ahead | 0.35 | 0.34 | 0.00 |
| 4 wk ahead | 0.32 | 0.30 | 0.02 |
| Season onset | 0.23 | 0.22 | 0.01 |
| Season peak percentage | 0.32 | 0.27 | 0.05 |
| Season peak week | 0.34 | 0.32 | 0.02 |

### Delayed case reporting impacts forecast score

In the seven seasons examined in this study, wILI percentages were often revised after first being reported. The frequency and magnitude of revisions varied by region. For example, in HHS Region 9, over 51% of initially reported wILI values ended up being revised by over 0.5 percentage points while in HHS Region 5 less than 1% of values were revised that much. Across all regions, 10% of observations were ultimately revised by more than percentage points.

When the first report of the wILI measurement for a given region-week was revised in subsequent weeks, we observed a corresponding strong negative impact on forecast accuracy. Larger revisions to the initially reported data were strongly associated with a decrease in the forecast score for the forecasts made using the initial, unrevised data. Specifically, among the four top-performing non-ensemble models (ReichLab-KCDE, LANL-DBM, Delphi-DeltaDensity1, and CU-EKF_SIRS), there was an average change in forecast score of -0.29 (95% CI: -0.39, -0.19) when the first observed wILI measurement was between 2.5 and 3.5 percentage points lower than the final observed value, adjusting for model, week-of-year, and target (Figure 5, see Methods for details on regression model). Additionally, we observed an expected change in forecast score of -0.24 (95% CI: -0.29, -0.19) when the first observed wILI measurement was between 1.5 and 2.5 percentage points higher than the final observed value. This pattern is similar for under-and over-reported values, although there were more extreme under-reported values than there were over-reported values. Finally, some of the variation in region-specific performance could be attributed to the frequency and magnitude of data revisions (data not shown).

**Figure 5:**
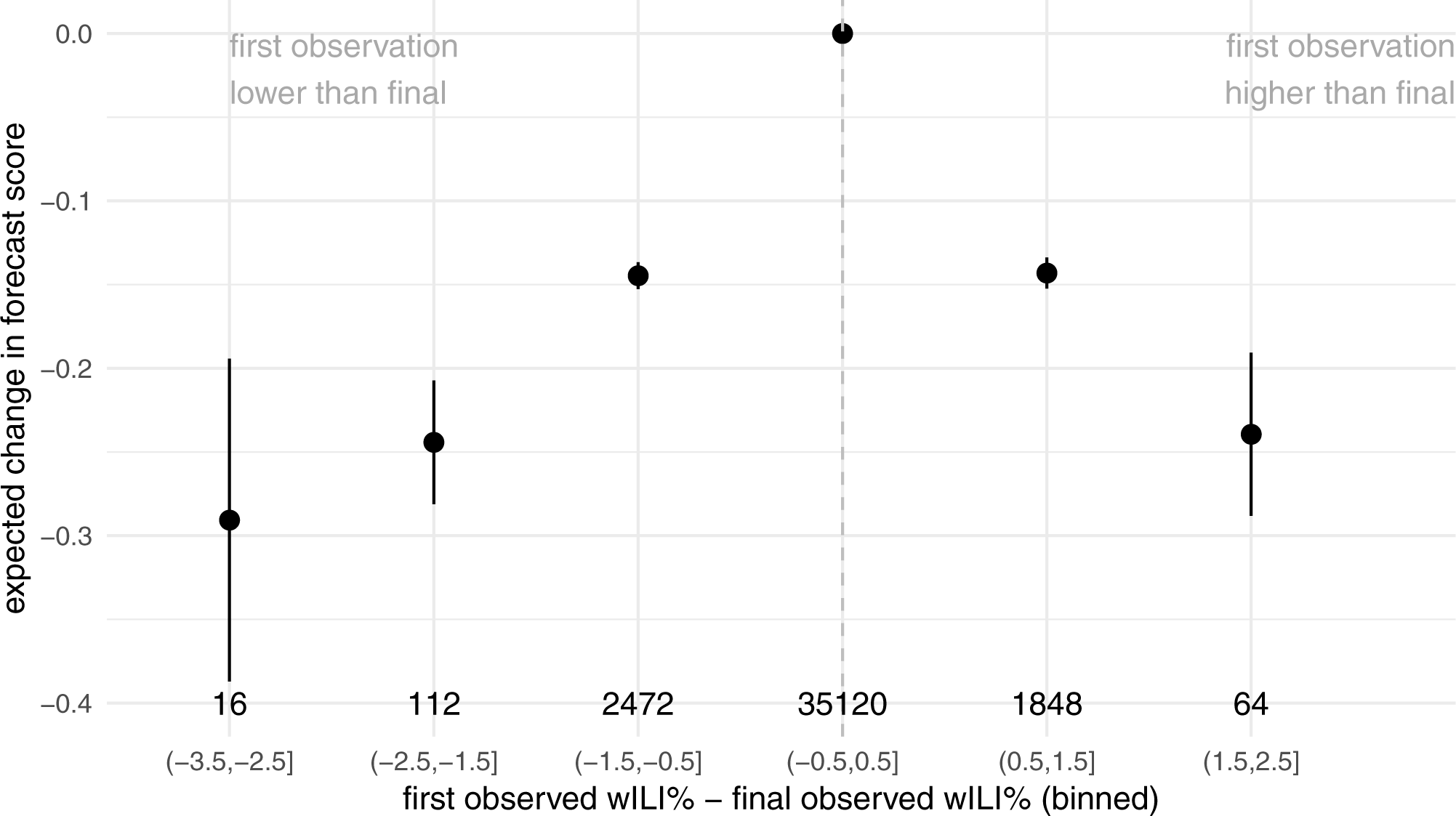
Model-estimated changes in forecast skill due to bias in initial reports of wILI %. The figure shows estimated coefficient values (and 95% confidence intervals) from a multivariable linear regression using model, week-of-year, target, and a categorized version of the bias in the first reported wILI % to predict forecast score. The x-axis labels show the range of bias (e.g. “(-0.5,0.5]” represents all observations whose first observations were within *±* 0.5 percentage points of the final reported value). Values to the left of the dashed grey line are observations whose first reported value were lower than the final. Y-axis values of less than zero (the reference category) represent decreases in expected forecast skill. The total number of observations in each category are shown above the x-axis labels.

## Discussion

This work presents the first large-scale comparison of real-time forecasting models from different modeling teams across multiple years. With the rapid increase in infectious disease forecasting efforts, it can be difficult to understand the relative importance of different methodological advances in the absence of an agreed-upon set of standard evaluations. We have built on the foundational work of CDC efforts to establish and evaluate models against a set of shared benchmarks which other models can use for comparison. Our collaborative, team science approach highlights the ability of multiple research groups working together to uncover patterns and trends of model performance that are harder to observe in single-team studies.

Seasonal influenza in the US, given the relative accessibility of historical surveillance data and recent history of coordinated forecasting ‘challenges’, is an important testbed system for understanding the current state of the art of infectious disease forecasting models. Using models from some of the most experienced forecasting teams in the country, this work reveals several key results about forecasting seasonal influenza in the US.

- A majority of models consistently showed higher accuracy than historical baseline forecasts, both in regions with more predictable seasonal trends and those with less consistent seasonal patterns (Figure 3B, D, E);
- A majority of the presented models showed consistent improvement over the historical baseline for 1 and 2 week-ahead forecasts, although fewer models consistently outperformed the baseline model for 3 and 4 week-ahead forecasts (Figure 3E);
- At the presented spatial and temporal resolutions for influenza forecasts, we did not identify substantial or consistent differences between high-performing models that rely on an underlying mechanistic (i.e., compartmental) model of disease transmission and those that are more statistical in nature (Table 2);
- Forecast accuracy is significantly degraded in some regions due to initial partially reported real-time data (Figure 5).

As knowledge and data about a given infectious disease system improve and become more granular, a common question among domain-area experts is whether mechanistic models will outperform more statistical approaches. However, the statistical vs. mechanistic model dichotomy is not always a clean distinction in practice. In the case of influenza, mechanistic models simulate a specific disease transmission process, whereas the data-generating mechanism for influenza-like illness captures much more than just disease transmission (e.g., clinical visitation behaviors, symptomatic diagnosis process, reporting process, a data-revision process, etc.). Therefore, a disease transmission model and the true underlying influenza-like illness data-generation process are fundamentally different, suggesting a limitation to purely mechanistic models in this context.

There are several important limitations to this work as presented. While we have assembled and analyzed a range of models from experienced influenza forecasting teams, there are large gaps in the types of data and models represented in our library of models. For example, relatively few additional data sources have been incorporated into these models, no models are included that explicitly incorporate information about circulating strains of influenza, and no model explicitly includes spatial relationships between regions. Given that several of the models rely on similar modeling frameworks, adding a more diverse set of modeling approaches would be a valuable contribution. Additionally, while seven seasons of forecasts from 22 models is the largest study we know of that compares models from multiple teams, this remains a less-than-ideal sample size to draw strong conclusions about model performance. Since each season represents a set of highly correlated dynamics across regions, there are not a lot of data from which to draw strong conclusions about comparative model performance.

What is the future of influenza forecasting in the US and globally? While long-run forecast accuracy for influenza will vary based on a variety of factors, including data quality and geographical scale, we expect to see continued forecast improvement through competition, collaboration, and methodological and technological innovation. We see particular promise in models that leverage different data sources, such as pathogen-specific and highly localized incidence data. Additionally, building ensemble models that capitalize on the strengths of a diverse set of individual component models will be critical to improving accuracy and consistency of models in all infectious disease forecasting settings. Ensemble forecasting was the motivation behind the creation of the FluSight Network and, while out of scope of this manuscript, will be the topic of future collaborative research for the group.

To advance infectious disease forecasting broadly, a complete enumeration and understanding of the challenges facing the field is critical. In this work, we have identified and quantified some of these challenges, specifically focusing on timely reporting of surveillance data. However, other barriers may be of equal or greater importance to continued improvement of forecasts. Often, researchers either lack access to or do not know how best to make use of novel data streams (e.g., Internet data, electronic medical health record data). Increased methodological innovation in models that merge together an understanding of biological drivers of disease transmission (e.g., strain-specific dynamics and vaccination effectiveness) with statistical approaches to combine data hierarchically at different spatial and temporal scales will be critical to moving this field forward. From a technological perspective, additional efforts to standardize data collection, format, storage, and access will increase interoperability between groups with different modeling expertise, improve accessibility of novel data streams, and continue to provide critical benchmarks and standards for the field.

Public health officials are still learning how to best integrate forecasts into real-time decision making. Close collaboration between public health policy-makers and quantitative modelers is necessary to ensure that forecasts have maximum impact and are appropriately communicated to the public and the broader public health community. Real-time implementation and testing of forecasting methods plays a central role in planning and assessing what targets should be forecasted for maximum public health impact.

## Methods

### FluSight Challenge Overview

Detailed methodology and results from previous FluSight challenges have been published[8, 10], and we summarize the key features of the challenge here.

During each influenza season, the wILI data are updated each week by the CDC. When the most recent data are released, the prior weeks’ reported wILI data may also be revised. The unrevised data, available at a particular moment in time, are available via the DELPHI real-time epidemiological data API beginning in the 2014/2015 season.[30] This API enables researchers to “turn back the clock” to a particular moment in time and use the data available at that time. This tool facilitates more accurate assessment of how models would have performed in real-time.

The FluSight challenges have defined seven forecasting targets of particular public health relevance. Three of these targets are fixed scalar values for a particular season: onset week, peak week, and peak intensity (i.e., the maximum observed wILI percentage). The remaining four targets are the observed wILI percentages in each of the subsequent four weeks (Figure 1B). A season has an onset week when at least three consecutive weeks are above a CDC-defined regional baseline for wILI. The first of these weeks is considered to be the onset week

The FluSight challenges have also required that all forecast submissions follow a particular format. A single submission file (a comma-separated text file) contains the forecast made for a particular epidemic week (EW) of a season. Standard CDC definitions of epidemic week are used.[31, 32, 33] Each file contains binned predictive distributions for seven specific targets across the 10 HHS regions of the US plus the national level. Each file contains over 8000 rows and typically is about 400KB in size.

To be included in the model comparison presented here, previous participants in the CDC FluSight challenge were invited to provide out-of-sample forecasts for the 2010/2011 through 2016/2017 seasons. For each season, files were submitted for epidemic week 40 (EW40) of the first calendar year of the season through EW20 of the following calendar year. (For seasons that contained an EW53, an additional file labeled EW53 was included.) For each model, this involved creating 233 separate forecast submission files, one for each of the weeks in the seven training seasons. In total, the forecasts represent over 40m rows and 2.5GB of data. Each forecast file represented a single submission file, as would be submitted to the CDC challenge. Each team created their submitted forecasts in a prospective, out-of-sample fashion, i.e., fitting or training the model only on data available before the time of the forecast (see Figure 1). All teams used the Delphi epidata API to retrieve ILINet data.[30] Some data sources (e.g. wILI data prior to the 2014/2015 season) were not archived in a way that made data reliably retrievable in this “real-time” manner. In these situations, teams were still allowed to use these data sources with best efforts made to ensure forecasts were made using only data available at the time forecasts would have been made.

### Summary of Models

Five teams each submitted between 1 and 9 separate models for evaluation (Table 1). A wide range of methodological approaches and modeling paradigms are included in the set of forecast models. For example, seven of the models utilize a compartmental structure (e.g. Susceptible-Infectious-Recovered), a model framework that explicitly encodes both the transmission and the susceptible-limiting dynamics of infectious disease outbreaks. Other less directly mechanistic models use statistical approaches to model the outbreak phenomenon by incorporating recent incidence and seasonal trends. One model, Delphi-Stat, is an ensemble model, a combination of other models from the Delphi team. Six models directly incorporate external data (i.e., not just the wILI measurements from the CDC ILINet dataset), including historical humidity data and Google search data. Three models stand out as being reference models. One shared feature of these models is that their forecasts do not depend on observed data from the season being forecasted. The Delphi-Uniform model always provides a forecast that assigns equal probability to all possible outcomes. The ReichLab-KDE model yields predictive distributions based entirely on data from other seasons using kernel density estimation (KDE) for seasonal targets and a generalized additive model with cyclic penalized splines for weekly incidence. The Delphi-EmpiricalTraj model uses KDE for all targets. The ‘historical baseline’ model named throughout the manuscript refers to the ReichLab-KDE model. Because this model represents a prediction that essentially summarizes historical data, we consider this model an appropriate baseline model to reflect historical trends. Once submitted to the central repository, the models were not updated or modified except in four cases to fix explicit bugs in the code that yielded numerical problems with the forecasts. (In all cases, the updates did not substantially change the performance of the updated models.) Re-fitting of models or tuning of model parameters was explicitly discouraged to avoid unintentional overfitting of models.

### Metric Used for Evaluation and Comparison

The log-score for a model *m* is defined as log *f*_*m*_(*z*^*∗*^*|***x**) where *f*_*m*_(*z|***x**) is the predicted density function from model *m* for target *Z* conditional on some data **x**, *z*^*∗*^ is the observed value of the target *Z*, and log is the natural logarithm. The log-score is a “proper” scoring rule, which has the practical implication that linear combinations (i.e., arithmetic means) of log scores will also be proper.[28]

Following CDC FluSight evaluation procedures, we computed modified log-scores for the targets on the wILI percentage scale such that predictions within *±* 0.5 percentage points are considered accurate, i.e., modified 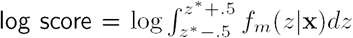. For the targets on the scale of epidemic weeks, predictions within *±* 1 week are considered accurate, i.e., modified 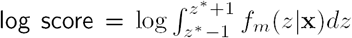. While this modification means that the resulting score is not formally a proper scoring rule, some have suggested that improper scores derived from proper scoring rules may, with large enough sample size, have negligible differences in practice.[28] Additionally, this modified log score has the advantage of having a clear interpretation and was motivated and designed by public health officials to reflect an accuracy of practical significance. Hereafter, we will refer to these modified log scores as simply log scores.

Average log scores can be used to compare models’ performance in forecasting for different locations, seasons, targets, or times of season. In practice, each model *m* has a set of log scores associated with it that are region-, target-, season-, and week-specific. We represent one specific scalar log score value as log *f*_*m,r,t,s,w*_(*z*^*∗*^*|***x**). These values can be averaged across any of the indices to create a summary measure of performance. For example,

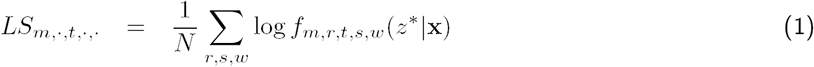

represents a log score for model *m* and target *t* averaged across all regions, seasons and weeks.

While log scores are not on a particularly interpretable scale, a simple transformation enhances interpretability substantially. Exponentiating an average log score yields a forecast score equivalent to the geometric mean of the probabilities assigned to the eventually observed outcome (or, more specifically for the modified log score, to regions of the distribution eventually considered accurate). The geometric mean is an alternative measure of central tendency to an arithmetic mean, representing the *N*^*th*^ root of a product of *N* numbers. Using the example from equation (1) above, we then have that

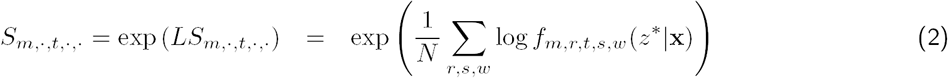

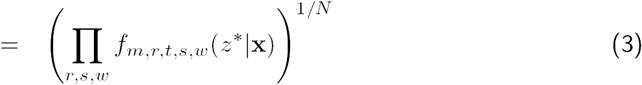

In this setting, this score *S* has the intuitive interpretation of being the average probability assigned to the true outcome (where average is considered to be a geometric average). Throughout the manuscript, we refer to an exponentiated average log score as an average score. In all cases, we compute the averages arithmetically on the log scale and only exponentiate before reporting and interpreting a final number. Therefore, all reported average scores can be interpreted as the corresponding geometric means, or as the correponding average probabilities assigned to the true outcome.

Following the convention of the CDC challenges, we only included certain weeks in the calculation of the average log scores for each target. This focuses model evaluation on periods of time that are more relevant for public health decision making. Forecasts of season onset are evaluated based on the forecasts that are received up to six weeks after the observed onset week within a given region. Peak week and peak intensity forecasts were scored for all weeks in a specific region-season up until the wILI measure drops below the regional baseline level for the final time. Week-ahead forecasts are evaluated using forecasts received four weeks prior to the onset week through forecasts received three weeks after the weighted ILI goes below the regional baseline for the final time. In a region-season without an onset, all weeks are scored. To ensure all calculated summary measures would be finite, all log scores with values of less than -10 were assigned the value -10, following CDC scoring conventions. All scores were based on “ground truth” values of wILI data obtained as of September 27, 2017.

### Specific model comparisons

#### Analysis of data revisions

The CDC publicly releases data on doctor’s office visits due to ILI each week. These data, especially for the most recent weeks, are occasionally revised, due to new or updated data being reported to the CDC since their last publication. While often these revisions are fairly minor or non-existent, at other times, these revisions can be substantial, changing the reported wILI value by over 50% of the originally reported value. Since the unrevised data are used by forecasters to generate current forecasts, real-time forecasts can be biased by the initially reported, preliminary data.

We used a regression model to analyze the impact of these unrevised reports on forecasting. Specifically, for each region and epidemic week we calculated the difference between the first and the last reported wILI values for each epidemic week for which forecasts were generated in the seven seasons under consideration. We then created a categorical variable (*X*) with a binned representation of these differences using the following six categories covering the entire range of observed values: (-3.5,-2.5], (-2.5,-1.5], …, (1.5,2.5]. Using the forecasting results from the four most accurate individual non-ensemble models, (ReichLab-KCDE, LANL-DBM, Delphi-DeltaDensity1, CU-EKF_SIRS), we then fit the following linear regression model

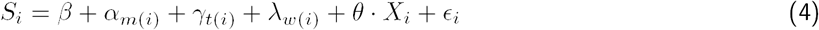

where *S*_*i*_ is the score, the index *i* indexes a specific set of subscripts *{m, r, t, s, w}*, and the *α*_*m*(*i*)_, *γ*_*t*(*i*)_, and *λ*_*w*(*i*)_ are model-, target-, and week-specific fixed effects, respectively. (The notation *m*(*i*) refers to the model contained in the *i*th observation of the dataset.) The error term is assumed to follow a Gaussian distribution with mean zero and an estimated variance parameter. The parameter of interest in the model is the vector *θ*, which represents the average change in score based on the magnitude of the bias in the latest available wILI value, adjusted for differences based on model, target, and week-of-season. The [*-*0.5, +0.5] change category was taken as a baseline category and the corresponding *θ* entry constrained to be 0, so that other *θ* entries represent deviations from this baseline.

#### Mechanistic vs. statistical models

There is not a consensus on a single best modeling approach or method for forecasting the dynamic patterns of infectious disease outbreaks in both endemic and emergent settings. Semantically, modelers and forecasters often use a dichotomy of mechanistic vs. statistical (or ‘phenomenological’) models to represent two different philosophical approaches to modeling. Mechanistic models for infectious disease consider the biological underpinnings of disease transmission, and in practice are implemented as variations on the Susceptible-Infectious-Recovered (SIR) model. Statistical models largely ignore the biological underpinnings and theory of disease transmission and focus instead on using data-driven, empirical and statistical approaches to make the best forecasts possible of a given dataset, or phenomenon.

However, in practice, this dichotomy is less clear than it is in theory. For example, statistical models for infectious disease counts may have an autoregressive term for incidence (e.g., as done by the ReichLab-SARIMA1 model). This could be interpreted as representing a transmission process from one time period to another. In another example, the LANL-DBM model has an explicit SIR compartmental model component but also utilizes a purely statistical model for the discrepancy of the compartmental model with observed trends. The models from Columbia University used a statistical ‘nowcasting’ approach for their 1-week ahead forecasts, but after that relied on different variations of an SIR model.

We categorized models according to whether or not they had any explicit compartmental framework (Table 1). We then took the top three performing compartmental models (i.e., models with some kind of an underlying compartmental structure) and compared their performance with the top three individual component models without compartmental structure. We excluded multi-model ensemble models (i.e., Delphi-Stat) from this comparison and also excluded the 1 week-ahead forecasts of the CU models from the compartmental model category, since they were generated by a statistical nowcast. Separately for each target, we compared the average score of the top three compartmental models to the average score of the top three non-compartmental models.

### Reproducibility and data availability

To maximize the reproducibility and data availability for this project, the data and code for the entire project (excluding specific model code) are publicly available. The project is available on GitHub[34], with a permanent repository stored on Zenodo[35]. All of the forecasts may be interactively browsed on the website http://flusightnetwork.io. A web applet with interactive visualizations of the model evaluations is available at https://reichlab.shinyapps.io/FSN-Model-Comparison/. Additionally, this manuscript was dynamically generated using R version 3.4.1 (2017-06-30), Sweave, and knitr, which are tools for intermingling manuscript text with R code that run the central analyses and minimize the chance for errors in transcribing or translating results.[36, 37].

## Acknowledgments and Disclaimers

The findings and conclusions in this report are those of the authors and do not necessarily represent the official position of the Centers for Disease Control and Prevention. JS and Columbia University disclose partial ownership of SK Analytics.

## References

[1] Natalie A. Molodecky, Isobel M. Blake, Kathleen M. O’Reilly, Mufti Zubair Wadood, Rana M. Safdar, Amy Wesolowski, Caroline O. Buckee, Ananda S. Bandyopadhyay, Hiromasa Okayasu, and Nicholas C. Grassly. Risk factors and short-term projections for serotype-1 poliomyelitis incidence in Pakistan: A spatiotemporal analysis. PLOS Medicine, 14(6):e1002323, jun 2017.

[2] Xiangjun Du, Aaron A King, Robert J Woods, and Mercedes Pascual. Evolution-informed forecasting of seasonal influenza A (H3N2). Science translational medicine, 9(413):eaan5325, oct 2017.

[3] Shweta Bansal, Gerardo Chowell, Lone Simonsen, Alessandro Vespignani, and Cécile Viboud. Big Data for Infectious Disease Surveillance and Modeling. Journal of Infectious Diseases, 214(Suppl 4):S375–S379, ec 2016.

[4] M.F. Myers, D.J. Rogers, J. Cox, A. Flahault, and S.I. Hay. Forecasting disease risk for increased epidemic preparedness in public health. Advances in Parasitology, 47:309–330, jan 2000.

[5] World Health Organization. Anticipating Emerging Infectious Disease Epidemics. Technical report, World Health Organization, Geneva, Switzerland, 2016.

[6] Jean-Paul Chretien, David Swedlow, Irene Eckstrand, Dylan George, Michael Johansson, Robert Huffman, and Andrew Hebbeler. Advancing Epidemic Prediction and Forecasting: A New US Government Initiative. Online Journal of Public Health Informatics, 7(1), 2015.

[7] Marc Lipsitch, Lyn Finelli, Richard T Heffernan, Gabriel M Leung, Stephen C Redd, and 2009 H1n1 Surveillance Group. Improving the evidence base for decision making during a pandemic: the example of 2009 influenza A/H1N1. Biosecurity and bioterrorism: biodefense strategy, practice, and science, 9(2):89–115, jun 2011.

[8] Matthew Biggerstaff, David Alper, Mark Dredze, Spencer Fox, Isaac Chun-Hai Fung, Kyle S. Hickmann, Bryan Lewis, Roni Rosenfeld, Jeffrey Shaman, Ming-Hsiang Tsou, Paola Velardi, Alessandro Vespignani, and Lyn Finelli. Results from the centers for disease control and prevention’s predict the 2013-2014 Influenza Season Challenge. BMC Infectious Diseases, 16(1):357, ec 2016.

[9] Morgan E Smith, Brajendra K Singh, Michael A Irvine, Wilma A Stolk, Swaminathan Subramanian, T Déirdre Hollingsworth, and Edwin Michael. Predicting lymphatic filariasis transmission and elimination dynamics using a multi-model ensemble framework. Epidemics, 18:16–28, 2017.

[10] Matthew Biggerstaff, Michael Johansson, David Alper, Logan C. Brooks, Prithwish Chakraborty, David C. Farrow, Sangwon Hyun, Sasikiran Kandula, Craig McGowan, Naren Ramakrishnan, Roni Rosenfeld, Jeffrey Shaman, Rob Tibshirani, Ryan J. Tibshirani, Alessandro Vespignani, Wan Yang, Qian Zhang, and Carrie Reed. Results from the second year of a collaborative effort to forecast influenza seasons in the United States. Epidemics, feb 2018.

[11] Cécile Viboud, Kaiyuan Sun, Robert Gaffey, Marco Ajelli, Laura Fumanelli, Stefano Merler, Qian Zhang, Gerardo Chowell, Lone Simonsen, and Alessandro Vespignani. The RAPIDD ebola forecasting challenge: Synthesis and lessons learnt. Epidemics, aug 2017.

[12] MA Rolfes, IM Foppa, S Garg, B Flannery, L Brammer, JA Singleton, Burns E, Jernigan D, C Reed, SJ Olsen, and J Bresee. Estimated influenza illnesses, medical visits, hospitalizations, and deaths averted by vaccination in the united states. https://www.cdc.gov/flu/about/disease/2015-16.htm.

[13] William W Thompson, David K Shay, Eric Weintraub, Lynnette Brammer, Nancy Cox, Larry J Anderson, and Keiji Fukuda. Mortality associated with influenza and respiratory syncytial virus in the united states. Jama, 289(2):179–186, 2003.

[14] Anice C Lowen, Samira Mubareka, John Steel, and Peter Palese. Influenza virus transmission is dependent on relative humidity and temperature. PLoS pathogens, 3(10):e151, 2007.

[15] Simon Cauchemez, Alain-Jacques Valleron, Pierre-Yves Boelle, Antoine Flahault, and Neil M Ferguson. Estimating the impact of school closure on influenza transmission from sentinel data. Nature, 452(7188):750, 2008.

[16] PhiResearchLab. Epidemic Prediction Initiative. https://predict.phiresearchlab.org/.

[17] Matthew Biggerstaff, Krista Kniss, Daniel B Jernigan, Lynnette Brammer, Joseph Bresee, Shikha Garg, Erin Burns, and Carrie Reed. Systematic assessment of multiple routine and near-real time indicators to classify the severity of influenza seasons and pandemics in the united states, 2003–04 through 2015–2016. American Journal of Epidemiology, 187:1040–1050, 2018.

[18] Sen Pei and Jeffrey Shaman. Counteracting structural errors in ensemble forecast of influenza outbreaks. Nature Communications, 8(1):925, ec 2017.

[19] Wan Yang, Alicia Karspeck, and Jeffrey Shaman. Comparison of Filtering Methods for the Modeling and Retrospective Forecasting of Influenza Epidemics. PLoS Computational Biology, 10(4):e1003583, apr 2014.

[20] Teresa K. Yamana, Sasikiran Kandula, and Jeffrey Shaman. Individual versus superensemble forecasts of seasonal influenza outbreaks in the United States. PLOS Computational Biology, 13(11):e1005801, nov 2017.

[21] Logan C Brooks, David C Farrow, Sangwon Hyun, Ryan J Tibshirani, and Roni Rosenfeld. epiforecast: Tools for forecasting semi-regular seasonal epidemic curves and similar time series. https://github.com/cmu-delphi/epiforecast-R, 2015.

[22] Logan C. Brooks, David C. Farrow, Sangwon Hyun, Ryan J. Tibshirani, and Roni Rosenfeld. Nonmechanistic forecasts of seasonal influenza with iterative one-week-ahead distributions. PLOS Computational Biology, 14(6):e1006134, jun 2018.

[23] Logan C. Brooks, David C. Farrow, Sangwon Hyun, Ryan J. Tibshirani, and Roni Rosenfeld. Flexible Modeling of Epidemics with an Empirical Bayes Framework. PLOS Computational Biology, 11(8):e1004382, aug 2015.

[24] Dave Osthus, James Gattiker, Reid Priedhorsky, and Sara Y. Del Valle. Dynamic Bayesian Influenza Forecasting in the United States with Hierarchical Discrepancy. arXiv, aug 2017.

[25] Evan L. Ray, Krzysztof Sakrejda, Stephen A. Lauer, Michael A. Johansson, and Nicholas G. Reich. Infectious disease prediction with kernel conditional density estimation. Statistics in Medicine, sep 2017.

[26] Evan L. Ray and Nicholas G. Reich. Prediction of infectious disease epidemics via weighted density ensembles. PLOS Computational Biology, 14(2):e1005910, feb 2018.

[27] George Sugihara and Robert M. May. Nonlinear forecasting as a way of distinguishing chaos from measurement error in time series. Nature, 344(6268):734–741, apr 1990.

[28] Tilmann Gneiting and Adrian E Raftery. Strictly proper scoring rules, prediction, and estimation. Journal of the American Statistical Association, 102(477):359–378, 2007.

[29] Sasikiran Kandula, Daniel Hsu, and Jeffrey Shaman. Subregional Nowcasts of Seasonal Influenza Using Search Trends. Journal of medical Internet research, 19(11):e370, nov 2017.

[30] DELPHI. Real-time Epidemiological Data API. https://github.com/cmu-delphi/delphi-epidata.

[31] New Mexico Department of Health. Indicator-Based Information System for Public Health Web. https://ibis.health.state.nm.us/resource/MMWRWeekCalendar.html.

[32] Jarad Niemi. MMWRweek: Convert Dates to MMWR Day, Week, and Year. https://CRAN.R-project.org/package=MMWRweek, 2015. R package version 0.1.1.

[33] Abhinav Tushar. pymmwr: MMWR weeks for Python. https://pypi.org/project/pymmwr/, 2018. python library version 0.2.2.

[34] A Tushar, NG Reich, T Yamana, D Osthus, C McGowan, EL Ray, SJ Fox, LC Brooks, and E Moore. FluSightNetwork: cdc-flusight-ensemble repository. https://github.com/FluSightNetwork/cdc-flusight-ensemble.

[35] A Tushar, NG Reich, T Yamana, D Osthus, C McGowan, EL Ray, SJ Fox, LC Brooks, and E Moore. FluSightNetwork/cdc-flusight-ensemble v1.0. https://doi.org/10.5281/zenodo.1255023.

[36] Yihui Xie. Dynamic Documents with R and knitr. Chapman and Hall/CRC, Boca Raton, Florida, 2nd edition, 2015. ISBN 978-1498716963.

[37] R Core Team. R: A language and environment for statistical computing. https://www.R-project.org/, 2017.

